# Advances in computer-assisted syndrome recognition and differentiation in a set of metabolic disorders

**DOI:** 10.1101/219394

**Authors:** Jean Tori Pantel, Max Zhao, Martin Atta Mensah, Nurulhuda Hajjir, Tzung-Chien Hsieh, Yair Hanani, Nicole Fleischer, Tom Kamphans, Stefan Mundlos, Yaron Gurovich, Peter M. Krawitz

## Abstract

Significant improvements in automated image analysis have been achieved over the recent years and tools are now increasingly being used in computer-assisted syndromology. However, the recognizability of the facial gestalt might depend on the syndrome and may also be confounded by severity of phenotype, size of available training sets, ethnicity, age, and sex. Therefore, benchmarking and comparing the performance of deep-learned classification processes is inherently difficult.

For a systematic analysis of these influencing factors we chose the lysosomal storage diseases Mucolipidosis as well as Mucopolysaccharidosis type I and II, that are known for their wide and overlapping phenotypic spectra. For a dysmorphic comparison we used Smith-Lemli-Opitz syndrome as a metabolic disease and Nicolaides-Baraitser syndrome as another disorder that is also characterized by coarse facies. A classifier that was trained on these five cohorts, comprising 288 patients in total, achieved a mean accuracy of 62%.

The performance of automated image analysis is not only significantly higher than randomly expected but also better than in previous approaches. In part this might be explained by our large training sets. We therefore set up a simulation pipeline that is suited to analyze the effect of different potential confounders, such as cohort size, age, sex, or ethnic background on the recognizability of phenotypes. We found that the true positive rate increases for all analyzed disorders for growing cohorts (*n*=[10…40]) while ethnicity and sex have no significant influence.

The dynamics of the accuracies strongly suggest that the maximum recognizability is a phenotype-specific value, that hasn’t been reached yet for any of the studied disorders. This should also be a motivation to further intensify data sharing efforts, as computer-assisted syndrome classification can still be improved by enlarging the available training sets.

Availability: software for classification: https://app.face2gene.com/research,

**Abbreviations**
DDx: Differential Diagnoses
DPDL: Deep Phenotyping for Deep Learning
DS: Down syndrome
FDNA: Facial Dysmorphology Novel Analysis
FNR: False Negative Rate
FPR: False Positive Rate
GAG: Glycosaminoglycan
HPO: Human Phenotype Ontology
LSD: Lysosomal Storage Disease
ML: Mucolipidosis
MPS I: Mucopolysaccharidosis type I
MPS II: Mucopolysaccharidosis type II
NCBRS: Nicolaides-Baraitser Syndrome
ROC: Receiver Operating Characteristics
SLOS: Smith-Lemli-Opitz Syndrome
TPR: True Positive Rate

## Introduction

In syndromology the information content of the facial gestalt is so extraordinarily high that several groups have worked on automated image analysis which assists in the diagnostic workup. Especially recent advances in computer vision improved pattern recognition on facial photos of syndromic patients (1-3). These approaches also have the potential to quantify the similarities of patients to any specific syndrome and to decide on a statistical basis whether two phenotypes are different (1, 4-7).

Face2Gene (FDNA Inc., Boston MA, USA) is a set of applications for pattern recognition in frontal photographs based on a neural network and crowd sourced medical data. It is available online (https://face2gene.com) and is used by a growing number of medical professionals. The CLINIC application of Face2Gene suggests differential diagnoses (DDx) for a patient based on frontal photos and phenotypic features annotated by the clinician. Whenever a diagnosis has been confirmed, the extracted de-identified information of the case is used to update the multiclass syndrome models and to improve predictions on similar cases in the future.

While Face2Gene CLINIC makes the latest classification models available that were trained on the entire set of suitable cases, a recently launched application, referred to as RESEARCH, that allows working with FDNA technology in a controllable environment. This app can be used to learn the facial gestalts of different cohorts that share for example disease-causing mutations in the same gene or pathway. The results of an experiment are gestalt models suitable for binary and multi-class comparisons. The true positive rates (TPRs) as well as the error rates of the multi-class problem are reported in a confusion matrix, whereas the pairwise comparison of cohorts are evaluated as receiver operating characteristics (ROC) curves.

Whenever the accuracies achieved by the gestalt models in the classification of photographs are higher than randomly expected, we conclude in this study that there is a recognizable facial pattern in a cohort. The possibility of a classifier to distinguish phenotypes is referred to as recognizability and if we quantify it, the phenotype of the controls matters as well. Especially when phenotypes of the same molecular subgroup, such as lysosomal storage diseases (LSDs), are studied, recognizability also means that a clinical entity can be delineated based on the facial gestalt. While this delineation of syndromic phenotypes has been reserved to few experts in the field, computer-assisted pattern recognition now objectifies this process and makes it even quantifiable.

However, it is self-explanatory that the extent of recognizability will not only depend on the “disease-phenotype” itself, but also on additional factors in the set-up of the studied cohorts. A systematic analysis of these potential confounders is a main aspect of this work.

## Patients and Methods

We analyzed the accuracy of gestalt models of the FDNA technology for a subset of disorders that have high phenotypic variability on the one hand and show a considerable phenotypic overlap on the other hand. We therefore selected Mucopolysaccharidoses (MPS I and II), which are inherited LSDs, resulting from mutations in enzymes catalyzing the breakdown of GAGs (8). Hardly any symptoms are present at birth, but appear usually during early childhood and progress in severity during adolescence (9). Most frequent features are abnormal bone size/shape including disproportionate short stature, contractures, intellectual disability and hearing loss (10). A coarse facial appearance, caused by progressive GAG deposition in the soft tissue, is of particular value for the establishment of the correct diagnosis (11). Important for our analysis was also that many cases of MPS have been published, due to its relatively high prevalence, and that the clinical diagnosis is easily confirmed by measuring enzyme activity or gene testing (12). MPS I, which is caused by mutations in alpha-L-iduronidase (*IDUA*), is clinically divided according to the severity of the symptoms into three subtypes: Hurler, Scheie, and Hurler-Scheie. MPS II, which is also referred to as Hunter syndrome, is caused by pathogenic mutations in the X-linked iduronate 2-sulfatase (*I2S*) and has high phenotypic similarity with MPS I. Both gene products are involved in similar processes in lysosomes where they catalyze the degradation of heparan sulfate and dermatan sulfate (11, 13). MPS I and II are still life-threatening diseases leading to a premature death and high morbidity (14, 15). However, for a few years enzyme replacement therapies are now available that have also been shown to attenuate the phenotypic features (16-19).

Another group of disease entities belonging to the lysosomal storage disorders are the different types of mucolipidosis (ML). In this work we focused on patients with pathogenic mutations in the alpha and beta subunits of N-acetyl-glucosamine-1-phosphotranseferase gene (*GNPTAB*). These subunits are part of a heterohexameric complex that catalyzes the first step of the synthesis of mannose-6-phosphat-recognition markers in the Golgi apparatus, which is a crucial step for correct trafficking of lysosomal enzymes (8).

### Hurler-like face

ML and MPS share common features, such as a coarsening of the face, organomegaly, skeletal malformations, and developmental delay, making them mutual DDx. Especially the facial gestalt of MPS and ML may be so similar that it is hard to tell the diseases apart without enzymatic or genetic testing, making them challenging choices for binary classification via computer vision.

We also added patients with Smith-Lemli-Opitz syndrome (SLOS) as another metabolic disorder and Nicolaides-Baraitser syndrome (NCBRS) as another disease phenotype that is also characterized by coarse facial features, in order to make the diagnostic workup more realistic. From a classification point of view, we are now dealing with a five-class problem.

The original sample set of 289 clinically diagnosed patients with MPS I, II, ML, SLOS, or NCBRS is based on case reports from the literature (for references see supplemental material). We conducted a literature search on PUBMED with the disease names and “case report” as search terms and looked for patient examples in textbooks. In addition to the patients’ photos, that were the basis for the automatized image analysis, we annotated the age, when the photo was taken and the presumed the ethnic background. For the analysis of the influence of ethnic background we also included the Down syndrome (DS) cases studied by Lumaka et al. If available, the disease-causing mutations were recorded in HGVS nomenclature and the phenotypic features were annotated in HPO terminology. A summary of the analyzed samples is shown in Figure 1. The entire case-based data collection is part of a larger knowledge base, called deep phenotyping for deep learning, DPDL, that can be accessed upon request and that serves as a set for benchmarking of computer-assisted image analysis.

**Figure 1.**
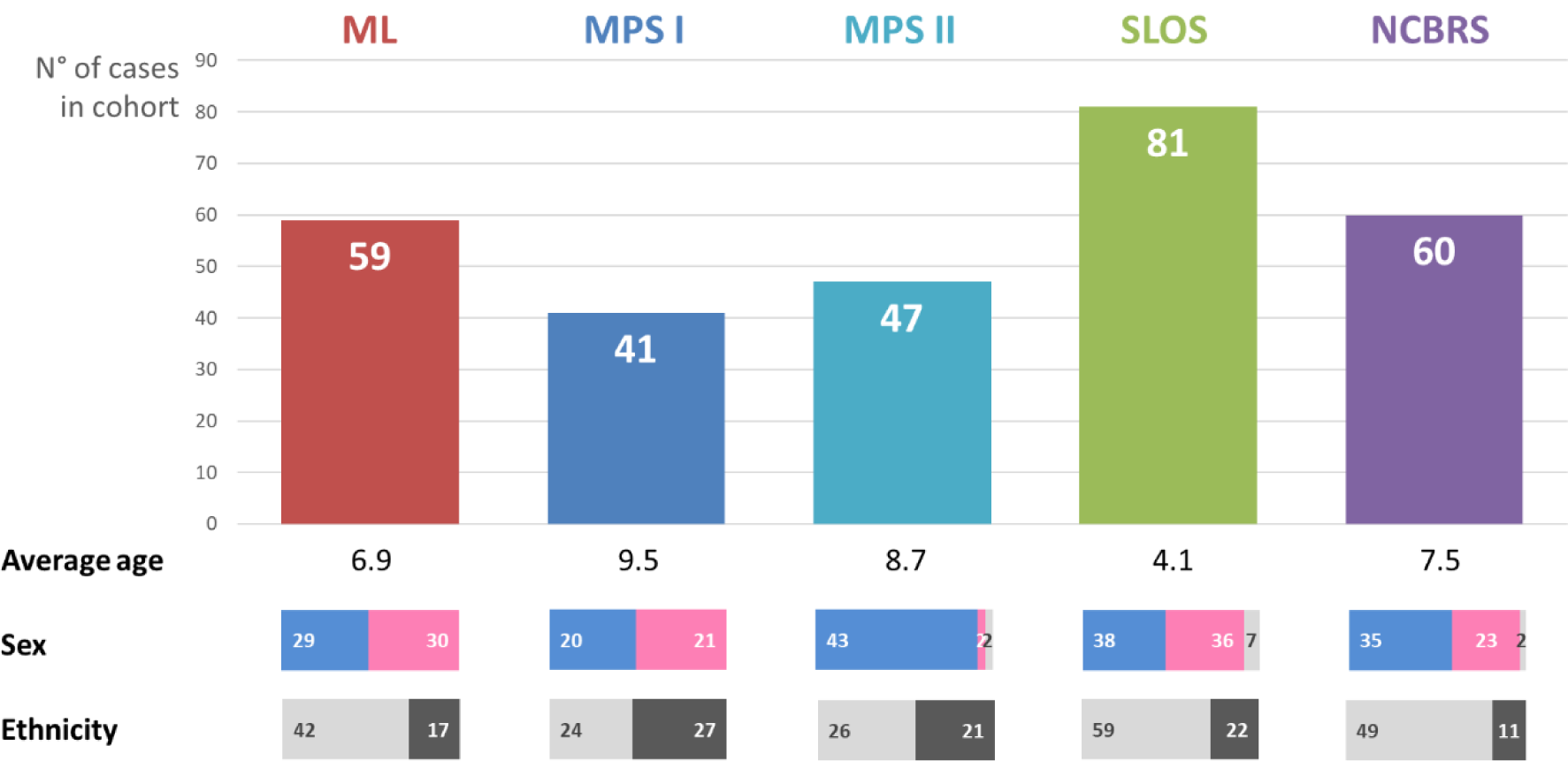
Overview of the original sample set with sex ratios (male/female/sex not mentioned) and ethnic backgrounds of European (left) vs. Non-European (right).

Cohorts for the phenotypic comparison were based on the clinical or molecular diagnosis and all experiments were run in Face2Gene’s RESEARCH application (version 17.6.2), which is accessible to its registered users.

The performance on subsets was evaluated by random down-sampling. A customized framework including the python requests library v2.18.4 was used to automatize the repetition of experiments and the TPRs of the resulting confusion matrices were averaged over 5 iterations for each setting. The scripts for the simulations are available on request and can be used to reproduce the results.

The potential cofounders of the technology that we choose to analyze in this work were cohort size, ethnic background and sex. The influence of the cohort size was analyzed by incrementing evenly sized subsets from 10 to 40. The change of the performance was fitted to a linear model and analyzed for significance using linregress of the SciPy library. The other potential cofounders ethnic background and sex were analyzed by excluding cohort size as a covariate. For these experiments, we sampled each cohort down to the greatest common size for each potential confounder. The greatest common size for the potential confounder male sex would e.g. be 20, because there are only 20 male patients with MPS I in our original sample set (see Figure 1). By this means cohort size has no influence on the performance and allowed an analysis of the potential confounders sex and ethnicity. Matthews correlation coefficient (MCC) is a measure of the quality of a two-class classification. Therefore, we reduced the multiclass confusion matrix to a two-class matrix for every diagnosis. Then we calculated the mean MCC for all iterations of the same experiment. If the difference of the MCCs of the potential confounder and control experiments was within the range of two standard deviations of the MCCs of the control experiments, we regarded the variable as not having a significant effect on the analyzed disease.

## Results

### Classification of the original sample set in Face2Gene CLINIC and RESEARCH

In Face2Gene CLINIC 30 possible differential diagnoses (DDx) are listed by default per case. If only a frontal facial photograph is uploaded and no further clinical features are annotated, these DDx represent syndromes that achieved the highest gestalt scores in the image analysis. Figure 2 shows the frequency of the five diagnoses in the study cohort among these 30 suggested diagnoses at the time of this study. MPS I and MPS II were combined in Face2Gene CLINIC under the phenotypic series of MPS. The right diagnosis was detected in most of the cases in all cohorts. As expected from the clinical description of the diseases, MPS and ML are both listed as possible differential diagnoses in more than half of the cases. SLOS and NCBRS occur in at least 13% of the cases as DDx among the 30 top-ranked diagnoses, which is also more than randomly expected given the fact that there are several hundreds of syndromes to choose from.

**Figure 2.**
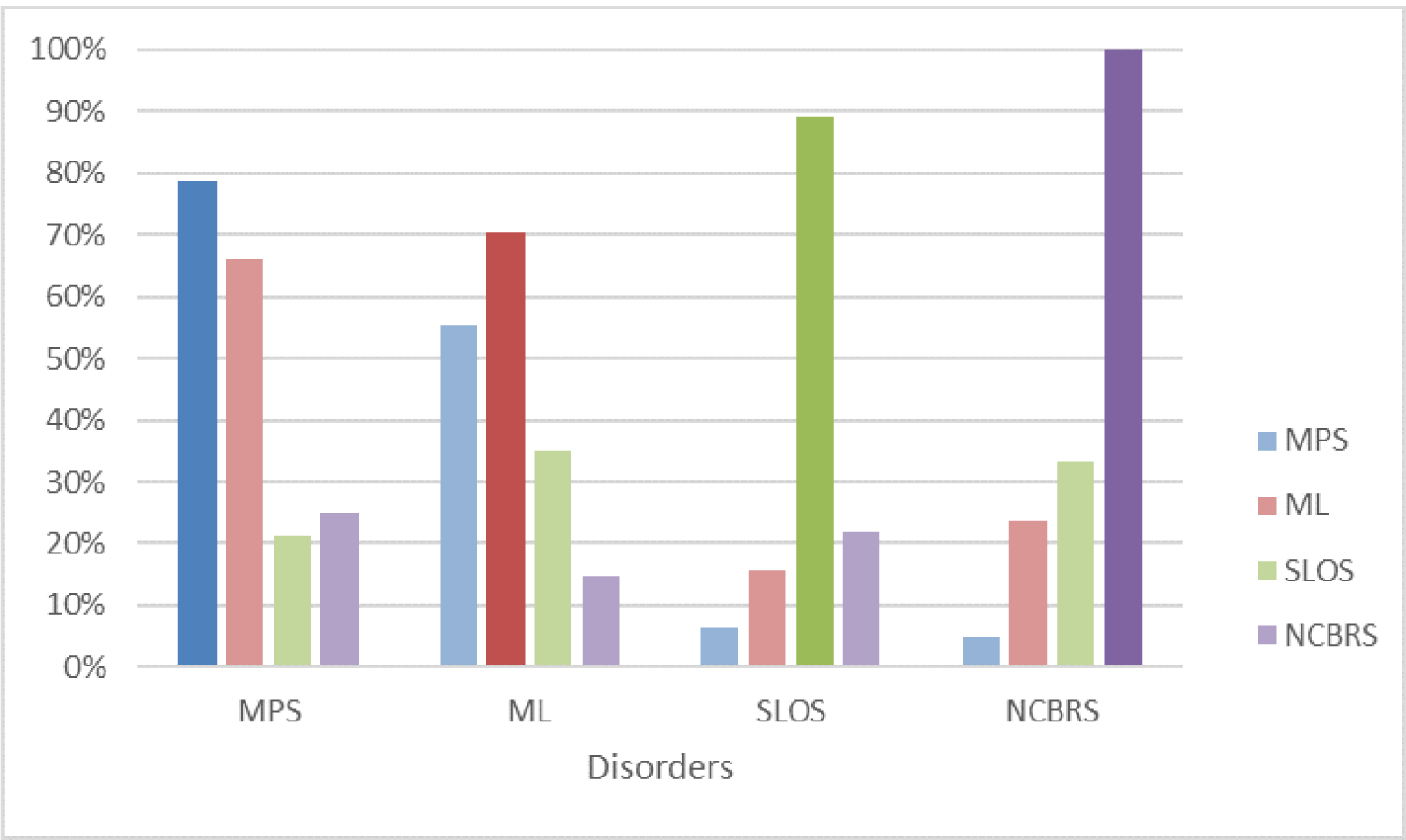
Frequency of occurence of the five disorders as differential diagnoses in the top 30 ranks in Face2Gene CLINIC, based on gestalt scores. MPS and ML often appear both as DDx, whereas they show up less frequently in SLOS and NCBRS.

While the patient data that was used for modeling the different phenotypes in Face2Gene CLINIC, is not directly accessible, Face2Gene RESEARCH allows *in silico* experiments with user defined cohorts. The resulting confusion matrix for the original sample set is shown in Figure 3 as a heat map. The stronger the field is colored, the higher is the TPR or FPR. The TPRs are higher than the FPRs for all cohorts, even for ML, MPS I and MPS II that are more frequently called as mutual false positives than as false positives of NCBRS or SLOS. Based on the similarity of the values in the columns, we calculated a dendrogram. The three lysosomal storage disorders MPS I, MPS II, and ML are more difficult to distinguish, as indicated by higher error rates in this cluster. Notably, the clustering of the five tested syndromes fits with their traditional clinical classification.

**Figure 3.**
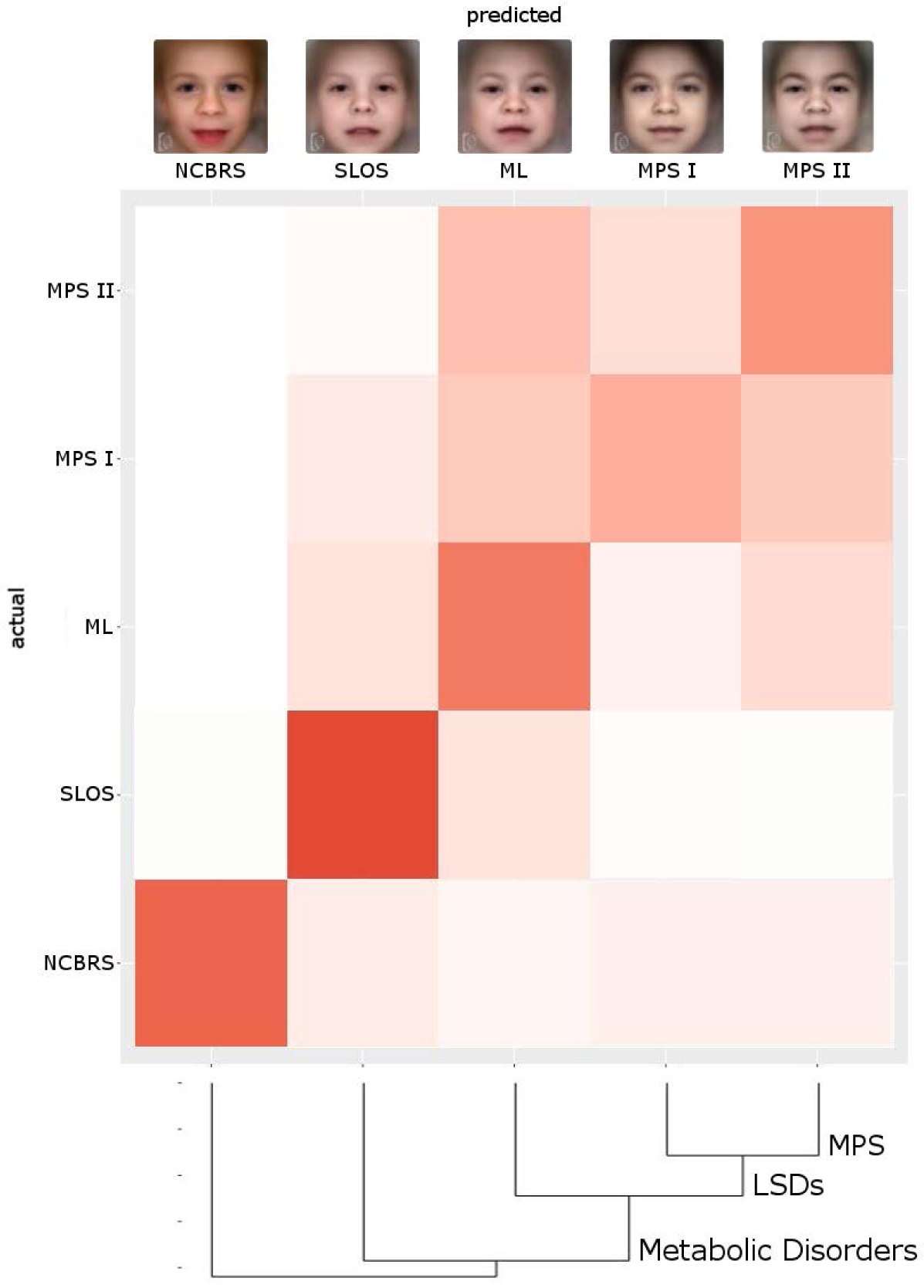
The performance of the gestalt-model in the multi-class problem in Face2Gene RESEARCH is shown as a color-coded confusion matrix, where deep red corresponds to a high value. True positive rates (TPR)are on the diagonal and false negatives and positives rates aside. The whole classification process achieves an accuracy of 62 %, which is significantly better than randomly expected (28 %). Photos on top show the average appearance of the disorder (mask) based on all cases of the respective cohort. The dendrogram is the result of the clustering analysis and visualizes the similarity of the disorders.

### Influence of growing cohort size on TPR

To analyze the influence of cohort size on the performance of the classification process, we increased the number of individuals per group stepwise from 10 to 40. TPRs with cohorts of only ten individuals were already higher than randomly expected and increased for all phenotypes with a growing cohort size, while the standard deviation decreased (Figure 4). The dynamics of the TPRs were fitted to a linear function and indicate that the full potential of computer-assisted classification hasn’t been reached yet with the available image data. However, we hypothesize that the number of images needed to reach the maximum recognizability could be different for each syndrome and might depend on the clinical variability of the phenotype. The TPRs of NCBRS and SLOS are the highest in comparison with the other cohorts. The lysosomal storage diseases are more frequently misclassified among each other than as SLOS or NCBRS. Notably, the MPS I-TPR nearly equals the fraction of MPS I cases falsely classified as MPS II., It is noteworthy, that ML is falsely classified to SLOS in around 14% and vice versa.

**Figure 4:**
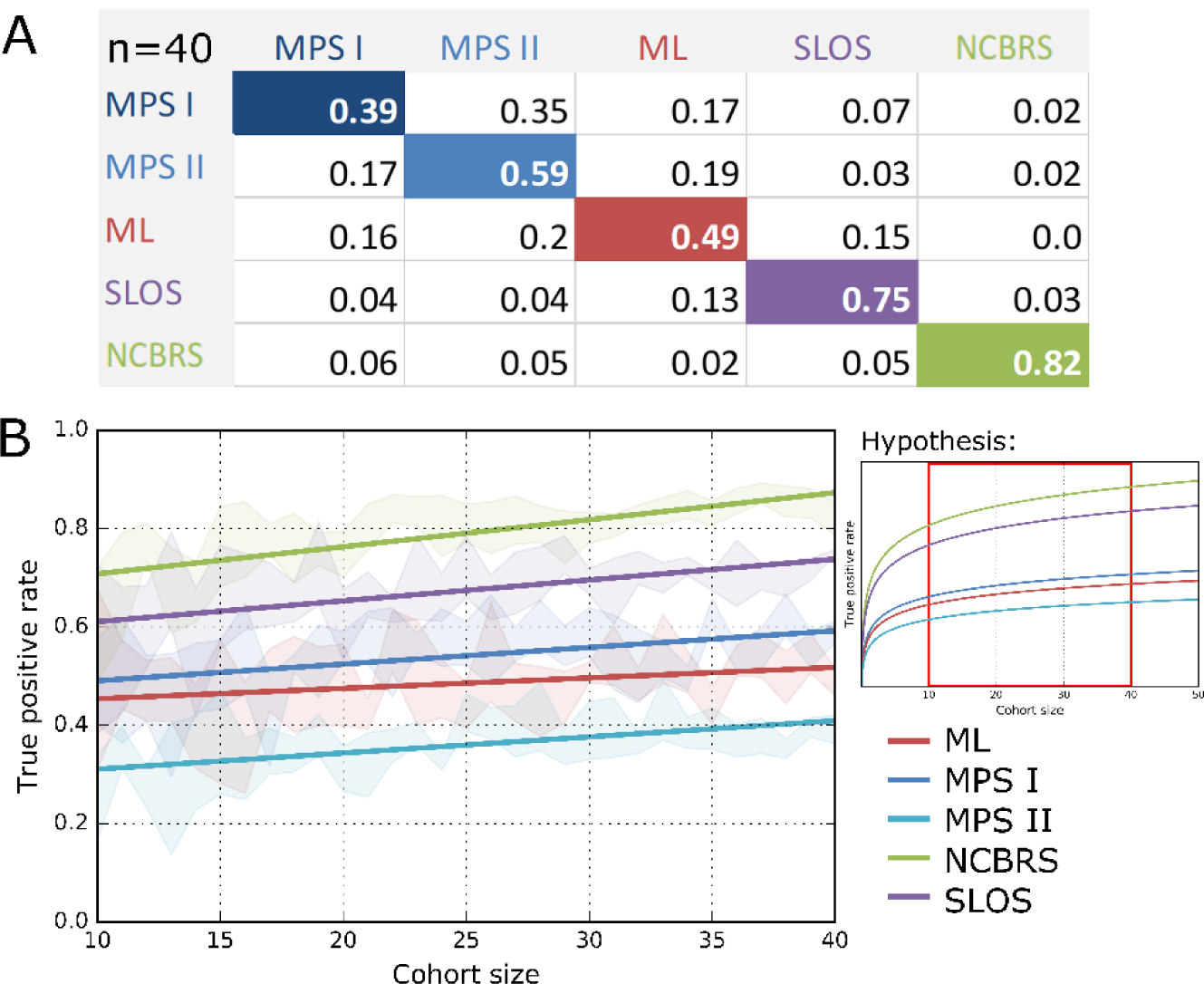
(A) Confusion matrix with TPRs and FPRs with a cohort size of n=40. (B) Course of TPRs with increasing cohort size with linear regression. The performance of the classification process was evaluated for equally sized cohorts from n=10 to n=40. The true positive rates for the prediction of the disorder improve with increasing cohort size and seem to approach different limits, indicating a difference in relative recognizability. Especially the prediction of SLOS and NCBRS benefit, when the classifier is trained on more cases. The inference of the correct lipid storage disorder increases less for larger cohort sizes.

### Effect of ethnic background or sex on performance

We hypothesized, that a bias in the set up of the cohorts with respect to the ethnic background or the sex, might affect the performance. In general, the performance should drop, if a true confounder is removed. If the performance increases instead after splitting up cohorts, this in an indicator that there is some characteristic feature, that can be more efficiently learned in a more homogeneous group of patients.

Lumaka *et al*. discussed in their study that certain features of Down syndrome, such as a deep nasal bridge and thick upper lips are less prominent in individuals of African descent. We adjusted for the same cohort size (n=19) and computed the MCCs that could be achieved in the classification of DS patients from Sub-Saharan Africa or Central Europe (table 1). Interestingly, we observed a substantially better performance for the DS model that was trained on the same ethnic background compared to a mixed setup of the cohorts (ΔMCC/STD for AFR vs. Mixed: 3.75 and ΔMCC/STD for CEU vs. Mixed: 2.70). This finding supports the hypothesis of a slightly different facial appearance of DS in Europeans and Africans.

**Table 1:**
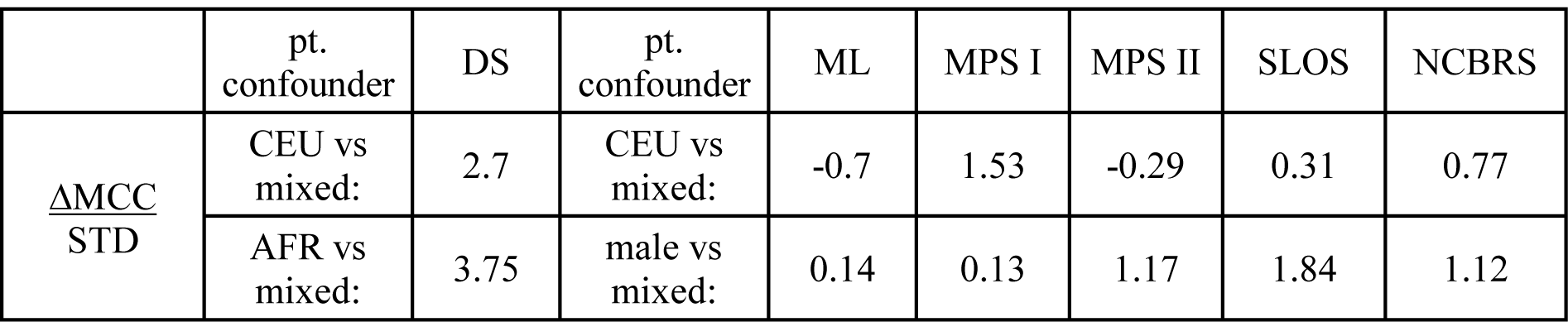
The classification of DS is more accurate on only European or African patients. These marked differences cannot be observed for ML, MPS I, MPSII, SLOS, and NCBRS. Also the restriction to only male patients has only a minor effect on the performance. The difference of MCCs for the binary classification of every disease was normalized by the standard deviations of MCCs that were computed in the mixed controls.

|  | pt.<br>confounder | DS | pt.<br>confounder | ML | MPS I | MPS II | SLOS | NCBRS |
| --- | --- | --- | --- | --- | --- | --- | --- | --- |
| $\frac{\Delta \text{MCC}}{\text{STD}}$ | CEU vs<br>mixed: | 2.7 | CEU vs<br>mixed: | -0.7 | 1.53 | -0.29 | 0.31 | 0.77 |
|  | AFR vs<br>mixed: | 3.75 | male vs<br>mixed: | 0.14 | 0.13 | 1.17 | 1.84 | 1.12 |

In contrast to DS, we did not observe such marked differences in the MCCs for MPS I, MPS II, ML, SLOS, and NCBRS, when running the experiments for n=22 cohorts that consisted only of European patients. An analysis for another background in these disorders was not possible due to a lack of sufficient patients.

Another potential confounder in the five-class problem of MPS I, MPS II, ML, SLOS, and NCBRS that we analyzed is sex. All but two of the MPS II patients were male, whereas the sex ratios for the other disorders were close to 1. To exclude cohort size as a covariate, we sampled down to n=20 in these experiments. Interestingly, the MCC for the MPS II classification did not decrease, when also all other cohorts consisted of male patients only. This indicates that a bias in the sex ratios does not affect the performance of the classification process substantially for the tested syndromes.

## Discussion

### General Recognizability

The TPRs that were achieved for all disorders in the five-class problems were higher than expected by random chance. Thus, our results show that the FDNA technology is capable of delineating gestalt differences even clinically similar phenotypes. This finding is especially remarkable for the phenotypes of MPS and ML and is also supported by high AUROC values in binary classifications (suppl. Fig. 1).

The difference in TPRs for the syndromes could be interpreted as different recognizabilities. Notably, SLOS and NCBRS are more recognizable than MPS I, MPS II, and ML. This corresponds to the results from the CLINIC app, where ML shows better ranks as a differential diagnosis for MPS and vice versa. These findings are in agreement with geneticist expert opinion, who label ML as highly similar to MPS.

The high TPRs found in our analyses corresponds to the results of two other studies on phenotypes of molecular pathway disorders. For Noonan syndrome as well as for GPI-anchor deficiencies, significant phenotypic substructures could be detected. This argues in favor of a more fine-grained phenotype modelling that might also be included in the CLINIC app prospectively.

### Ethnicity

MPS I could be more effectively differentiated from MPS II when cohorts were restricted to the same ethnic background. A possible explanation for this slight increase in performance might be, that there are certain features that are restricted or more prominent in European patients and that might therefore be learned more effectively if more cases are used for training the model. This issue has already been discussed for other disorders such as Fragile-X syndrome and Down syndrome were ethnic specific differences in the feature presentation are known (7, 20). Although we could replicate these effects for DS, we did not see a prominent change in the performance in the other phenotypes, which indicates that that ethnic background is not a strong confounder in the classification process.

### Sex

The human face shows a sexual dimorphism, possibly even at an early age, making sex a potential confounder in any facial image analysis process (21). The classification accuracies in our experiments that were based on data sets adjusted to individuals of the same sex, did not significantly differ, suggesting that the classification method is robust to sex as a confounder. Also the mean MCCs showed no significant change when training the classifier on only male individuals as compared to a training cohort consisting of both sexes. Our interpretation is that sex does not confound the classification of MPS I, MPS II, ML, SLOS, and NCBRS.

### Benchmarking

We are just beginning to understand the potential of computer assisted image analysis in the field of syndromology. In this work we have presented a general approach to study the recognizability of a phenotype and to test the confounding effect of variables such as ethnicity or sex. We have applied this framework to a selection of metabolic disorders, however, it is applicable to any other disorders.

It would be most interesting to compare the performance of the FDNA technology to the accuracies of other, previously published approaches of an automated image analysis of syndromic patients. Comparative evaluation, however, is impeded by the lack of a publicly available data set for benchmarking. Earlier benchmarking approaches merely relied on the comparison to a human classification performance. To achieve an objective evaluation computer vision, we strongly advocate to build a resource for image data of molecularly confirmed syndromic cases.

## References

1. Basel-Vanagaite L, Wolf L, Orin M, Larizza L, Gervasini C, Krantz ID, et al. Recognition of the Cornelia de Lange syndrome phenotype with facial dysmorphology novel analysis. Clin Genet. 2016;89(5):557–63.

2. Ferry Q, Steinberg J, Webber C, FitzPatrick DR, Ponting CP, Zisserman A, et al. Diagnostically relevant facial gestalt information from ordinary photos. Elife. 2014;3:e02020.

3. Loos HS, Wieczorek D, Wurtz RP, von der Malsburg C, Horsthemke B. Computer-based recognition of dysmorphic faces. Eur J Hum Genet. 2003;11(8):555–60.

4. Boehringer S, Vollmar T, Tasse C, Wurtz RP, Gillessen-Kaesbach G, Horsthemke B, et al. Syndrome identification based on 2D analysis software. Eur J Hum Genet. 2006;14(10):1082–9.

5. Vollmar T, Maus B, Wurtz RP, Gillessen-Kaesbach G, Horsthemke B, Wieczorek D, et al. Impact of geometry and viewing angle on classification accuracy of 2D based analysis of dysmorphic faces. Eur J Med Genet. 2008;51(1):44–53.

6. Gripp KW, Baker L, Telegrafi A, Monaghan KG. The role of objective facial analysis using FDNA in making diagnoses following whole exome analysis. Report of two patients with mutations in the BAF complex genes. Am J Med Genet A. 2016;170(7):1754–62.

7. Lumaka A, Cosemans N, Lulebo Mampasi A, Mubungu G, Mvuama N, Lubala T, et al. Facial dysmorphism is influenced by ethnic background of the patient and of the evaluator. Clin Genet. 2017;92(2):166–71.

8. Fenzl CR, Teramoto K, Moshirfar M. Ocular manifestations and management recommendations of lysosomal storage disorders I: mucopolysaccharidoses. Clin Ophthalmol. 2015;9:1633–44.

9. Muenzer J. Overview of the mucopolysaccharidoses. Rheumatology (Oxford). 2011;50 Suppl 5:v4–12.

10. Zschocke J, Hoffmann GF, Burlina AB, Milupa AG. Vademecum metabolicum: diagnosis and treatment of inborn errors of metabolism. 3rd rev. ed. Friedrichsdorf Stuttgart: Milupa Metabolics; Schattauer; 2011. x, 174 p. p.

11. Scarpa M. Mucopolysaccharidosis Type II. Adam MP, Ardinger HH, Pagon RA, et al, editors GeneReviews^®^ [Internet]. 2007.

12. Baehner F, Schmiedeskamp C, Krummenauer F, Miebach E, Bajbouj M, Whybra C, et al. Cumulative incidence rates of the mucopolysaccharidoses in Germany. J Inherit Metab Dis. 2005;28(6):1011–7.

13. Scott HS, Bunge S, Gal A, Clarke LA, Morris CP, Hopwood JJ. Molecular genetics of mucopolysaccharidosis type I: diagnostic, clinical, and biological implications. Hum Mutat. 1995;6(4):288–302.

14. Jones SA, Almassy Z, Beck M, Burt K, Clarke JT, Giugliani R, et al. Mortality and cause of death in mucopolysaccharidosis type II-a historical review based on data from the Hunter Outcome Survey (HOS). J Inherit Metab Dis. 2009;32(4):534–43.

15. Hendriksz CJ, Berger KI, Lampe C, Kircher SG, Orchard PJ, Southall R, et al. Health-related quality of life in mucopolysaccharidosis: looking beyond biomedical issues. Orphanet J Rare Dis. 2016;11(1):119.

16. Kubaski F, Yabe H, Suzuki Y, Seto T, Hamazaki T, Mason RW, et al. Hematopoietic Stem Cell Transplantation for Patients with Mucopolysaccharidosis II. Biol Blood Marrow Transplant. 2017.

17. Bradley LA, Haddow HRM, Palomaki GE. Treatment of mucopolysaccharidosis type II (Hunter syndrome): results from a systematic evidence review. Genet Med. 2017.

18. Rodgers NJ, Kaizer AM, Miller WP, Rudser KD, Orchard PJ, Braunlin EA. Mortality after hematopoietic stem cell transplantation for severe mucopolysaccharidosis type I: the 30-year University of Minnesota experience. J Inherit Metab Dis. 2017;40(2):271–80.

19. Watson HA, Holley RJ, Langford-Smith KJ, Wilkinson FL, van Kuppevelt TH, Wynn RF, et al. Heparan sulfate inhibits hematopoietic stem and progenitor cell migration and engraftment in mucopolysaccharidosis I. J Biol Chem. 2014;289(52):36194–203.

20. Schwartz CE, Phelan MC, Pulliam LH, Wilkes G, Vanner LV, Albiez KL, et al. Fragile X syndrome: incidence, clinical and cytogenetic findings in the black and white populations of South Carolina. Am J Med Genet. 1988;30(1-2):641–54.

21. Zhang W, Smith ML, Smith LN, Farooq A. Gender recognition from facial images: two or three dimensions? J Opt Soc Am A Opt Image Sci Vis. 2016;33(3):333–44.

